# Beyond 2/3 and 1/3: the complex signatures of sex-biased admixture on the X chromosome

**DOI:** 10.1101/016543

**Authors:** Amy Goldberg, Noah A Rosenberg

**Affiliations:** Department of Biology, Stanford University, Stanford, CA, 94305-5020 USA

## Abstract

Sex-biased demography, in which parameters governing migration and population size differ between females and males, has been studied through comparisons of X chromosomes, which are inherited sex-specifically, and autosomes, which are not. A common form of sex bias in humans is sex-biased admixture, in which at least one of the source populations differs in its proportions of females and males contributing to an admixed population. Studies of sex-biased admixture often examine the mean ancestry for markers on the X chromosome in relation to the autosomes. A simple framework noting that in a population with equally many females and males, 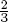 of X chromosomes appear in females, suggests that the mean X-chromosomal admixture fraction is a linear combination of female and male admixture parameters, with coefficients 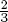 and 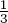, respectively. Extending a mechanistic admixture model to accommodate the X chromosome, we demonstrate that this prediction is not generally true in admixture models, though it holds in the limit for an admixture process occurring as a single event. For a model with constant ongoing admixture, we determine the mean X-chromosomal admixture, comparing admixture on female and male X chromosomes to corresponding autosomal values. Surprisingly, in reanalyzing African-American genetic data to estimate sex-specific contributions from African and European sources, we find that the range of contributions compatible with the excess African ancestry on the X chromosome compared to autosomes has a wide spread, permitting scenarios either without male-biased contributions from Europe or without female-biased contributions from Africa.

## Introduction

Comparisons of the X chromosome and the autosomes provide a strategy for understanding the history of sex-biased demography [1–14]. Unlike the autosomes, the X chromosome follows a sex-specific inheritance pattern, with females inheriting two copies, one from the mother and one from the father, and males inheriting only a single copy from the mother. As a consequence, demographic differences between females and males—in such phenomena as the breeding population size, the variance of reproductive success, and migration rates—can be studied by examining differences in patterns of genetic variation between X chromosomes and autosomes.

Many of the best-known cases of sex-biased patterns in human demography involve recently admixed populations [15–31]. During the formation of such populations, sex-biased admixture occurs if one or more of the source populations contributes different fractions of the females and males to the admixed group. What patterns are expected for X chromosomes and autosomes in an admixed population formed through a sex-biased admixture process? An initial hypothesis, reflecting the fact that in a population with equally many females and males, 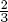 of the X chromosomes are in females and 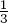 are in males, is that if 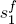 is the fraction of females originating from population 1 and 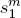 is the fraction of males originating from population 1, then the fraction of ancestry from population 1 for a site on the X chromosome is [19]

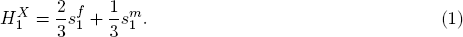

This simple linear combination has been used to estimate the sex-specific contributions from females and males to African-American and Latino populations [19, 31]. As we will show, however, it presumes a very specific history for the admixture process, a history often not viewed as reasonable for practical admixture scenarios.

Here, we extend a two-sex mechanistic admixture model [32] to incorporate the genetic signatures of sex-specific admixture patterns on the X chromosome. We derive a recursive expression for the expectation of the X-chromosomal admixture fraction as a function of sex-specific admixture parameters, demonstrating that the X-chromosomal admixture is obtained from a more complex formula than in the simple 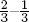 weighting (eq. (1)). The limiting mean X-chromosomal admixture is a predictable function of female and male contributions from the source populations, but among cases we consider, only has the 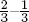 weighting for a single admixture event that takes place at a single point in time. During the approach to this limit, the behavior of the mean X-chromosomal admixture is dependent on the time since admixture. For a single admixture event and for constant ongoing admixture, we characterize the difference between the limit and the mean X-chromosomal admixture under the model as a function of time. We reinterpret admixture patterns in recently admixed African Americans, demonstrating that consideration of both the admixture model and the time since admixture is important for estimating sex-specific admixture contributions.

## Materials and Methods

### A mechanistic, sex-specific model for admixture histories

We follow the discrete-time model and notation of Verdu & Rosenberg [33] and Goldberg et al. [32], in which two source populations, *S*_1_ and *S*_2_, contribute to an admixed population, *H* (Fig. 1). For each population, female and male contributions are considered separately. At generation *g*, the contribution of sex *δ*, with *δ* ∈ {*f, m*}, from source *S*_*α*_, with *α* ∈ {1, 2}, is 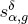. Corresponding contributions from *H* are denoted 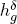. Thus, for example, 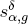 denotes the fraction in generation *g* of individuals of sex *δ* originating in the previous generation in population *S*_*α*_. The female and male contributions from *S*_1_, *H*, and *S*_2_ at generation *g* are 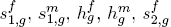, and 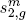.

**Figure 1:**
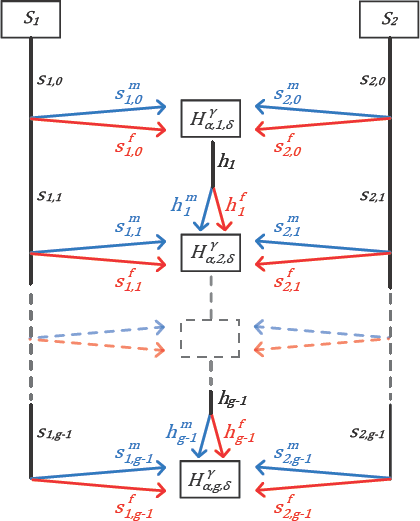
Sex-specific mechanistic model of admixture. At each generation *g*, females and males from each of two source populations, *S*_1_ and *S*_2_, contribute to the admixed population, *H*. Contributions can vary in time. The fraction of admixture from source population α ∈ {1, 2} for chromosomal type γ ∈ {*A, X*} in an admixed individual of sex δ ∈ {*f, m*} is 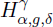. This model, by considering different chromosomal types, generalizes the model of Goldberg et al. [32].

We recall key relations among the sex-specific parameters [32, eqs. 1–6]. First, the total contribution from each population is the mean of female and male contributions: 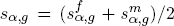 and 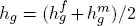. Also, as each parameter is a probability, the total female and male contributions separately sum to one: 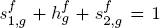 and 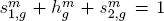. For *g* = 0, the admixed population does not yet exist, so that 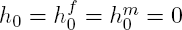 and 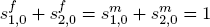.

Under this general framework, we define sex bias as a difference in the contributions from one or more source populations to the admixed population. That is, an admixture process is considered sex-biased if 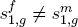 or 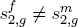 or both. Thus, sex bias involves differences between females and males entering from a source population, rather than a difference between parameters from the two source populations; after the founding, sex bias can occur in one source but not the other.

Considering the three populations (*S*_1_, *H, S*_2_) from which the parents of an individual from the admixed population can originate, we have nine possible ordered parental pairings (Table 1). As in Goldberg et al. [32], we study the random variable representing the admixture fraction of a random individual in the admixed population. Whereas Goldberg et al. [32] examined admixture only for the autosomes, here we also study the X chromosome. We define 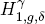 as the admixture fraction of chromosomal type *γ* sampled in individuals of sex *δ* from the admixed population *H* at generation *g*. Thus, 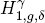 is the probability that a random site on an autosome (*γ* = *A*) or the X chromosome (*γ* = *X*) in a random individual of sex *δ* in *H* in generation *g* ultimately traces to *S*_1_.

**Table 1:** Recursion for the X-chromosomal and autosomal admixture fractions of a randomly chosen female or male from an admixed population at generation *g*, given a set of parents. With two source populations and the contributions from the admixed population to itself, a random individual has nine possible parental pairings, for each of which the probability is listed.

| Case | Female parent's population | Male parent's population | Probability | Admixture in males |  | Admixture in females |  |
| --- | --- | --- | --- | --- | --- | --- | --- |
| | | | | $H_{1,g,m}^A$ | $H_{1,g,m}^X$ | $H_{1,g,f}^A$ | $H_{1,g,f}^X$ |
| 1 | $S_1$ | $S_1$ | $s_{1,g-1}^f s_{1,g-1}^m$ | 1 | 1 | 1 | 1 |
| 2 | $S_1$ | $H$ | $s_{1,g-1}^f h_{g-1}^m$ | $\frac{1 + H_{1,g-1,m}^A}{2}$ | 1 | $\frac{1 + H_{1,g-1,m}^A}{2}$ | $\frac{1 + H_{1,g-1,m}^X}{2}$ |
| 3 | $S_1$ | $S_2$ | $s_{1,g-1}^f s_{2,g-1}^m$ | $\frac{1}{2}$ | 1 | $\frac{1}{2}$ | $\frac{1}{2}$ |
| 4 | $H$ | $S_1$ | $h_{g-1}^f s_{1,g-1}^m$ | $\frac{1 + H_{1,g-1,f}^A}{2}$ | $H_{1,g-1,f}^X$ | $\frac{1 + H_{1,g-1,f}^A}{2}$ | $\frac{1 + H_{1,g-1,f}^X}{2}$ |
| 5 | $H$ | $H$ | $h_{g-1}^f h_{g-1}^m$ | $\frac{H_{1,g-1,f}^A + H_{1,g-1,m}^A}{2}$ | $H_{1,g-1,f}^X$ | $\frac{H_{1,g-1,f}^A + H_{1,g-1,m}^A}{2}$ | $\frac{H_{1,g-1,f}^X + H_{1,g-1,m}^X}{2}$ |
| 6 | $H$ | $S_2$ | $h_{g-1}^f s_{2,g-1}^m$ | $\frac{H_{1,g-1,f}^A}{2}$ | $H_{1,g-1,f}^X$ | $\frac{H_{1,g-1,f}^A}{2}$ | $\frac{H_{1,g-1,f}^X}{2}$ |
| 7 | $S_2$ | $S_1$ | $s_{2,g-1}^f s_{1,g-1}^m$ | $\frac{1}{2}$ | 0 | $\frac{1}{2}$ | $\frac{1}{2}$ |
| 8 | $S_2$ | $H$ | $s_{2,g-1}^f h_{g-1}^m$ | $\frac{H_{1,g-1,m}^A}{2}$ | 0 | $\frac{H_{1,g-1,m}^A}{2}$ | $\frac{H_{1,g-1,m}^X}{2}$ |
| 9 | $S_2$ | $S_2$ | $s_{2,g-1}^f s_{2,g-1}^m$ | 0 | 0 | 0 | 0 |

In contrast to autosomal DNA, for which both female and male offspring receive a single copy of each chromosome from each parent, the X chromosome is inherited sex-specifically. That is, female offspring inherit one copy of the X chromosome from the mother and one from the father, but males inherit only a maternal copy. Therefore, whereas the autosomal admixture fractions sampled in females and males from the admixed population, 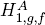 and 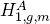, are identically distributed, the female and male X-chromosomal admixture fractions, 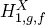 and 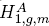, have different distributions. As a result, whereas Goldberg et al. [32] could uncouple 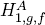 and 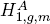 and consider recursions for these two quantities separately (both were identical and only one needed to be studied), here we examine the distributions of 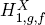 and 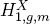 as a coupled pair of recursions.

Table 1 reports the probabilities that a random individual of a given sex from the admixed population has each possible set of parents, *l*, as well as the admixture fraction of the individual for source population 1 conditional on the parental pairing. The table provides a basis for recursively computing the fraction of admixture from *S*_1_ for a random individual of sex *δ* from the admixed population at generation *g*, 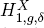. For *g* = 1, the founding event of the admixed population, the admixed population does not yet exist, so only cases 1, 3, 7, and 9, the four cases in which both parents are from the source populations, are considered.

Conditional on the previous generation, 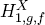 and 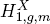 are independent random variables. The female X-chromosomal admixture fraction depends on both the female and male admixture fractions in the previous generation, 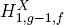 and 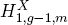, but the male X-chromosomal admixture fraction depends only on 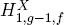. Therefore, the female X-chromosomal admixture fraction can equivalently be written as a function of the female X-chromosomal admixture fractions in the previous generation, 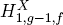, and from two generations ago, 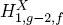.

### Expectation of the X-chromosomal fraction of admixture

Following Verdu & Rosenberg [33] and Goldberg et al. [32], we can find the expectation of the X-chromosomal admixture fraction of an individual of specified sex randomly chosen from the admixed population. Using the law of total expectation to consider the random parental pairing *L*, we sum over the nine possible pairings *l* (Table 1). For the mean X-chromosomal admixture fraction of a randomly chosen admixed individual of sex *δ* sampled at generation *g*, we have

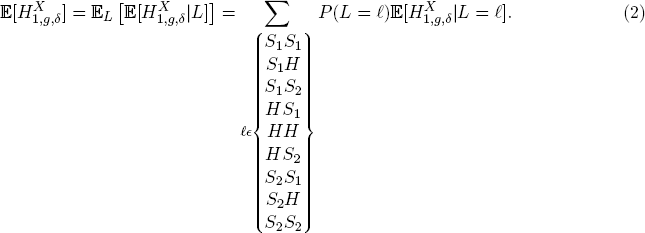

Applying eq. (2) using Table 1 to consider X chromosomes sampled in females and males from the admixed population, for the first generation, the initial condition is

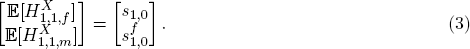

For *g* ≥ 2, we have

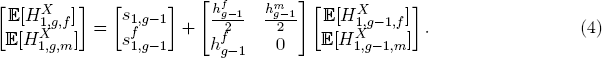

For the autosomes, for each sex *δ*, the expectation of the admixture fraction depends only on the total (non-sex-specific) contributions from the source populations and the corresponding expectation in the previous generation [32, eq. 19],

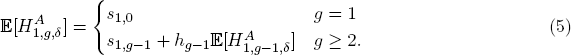

By contrast, the expectations of the female and male X-chromosomal admixture fractions depend also on the sex-specific contributions in the previous generation (eq. (4)).

Because the inheritance pattern is identical for X chromosomes and autosomes in females, the mean X-chromosomal admixture in females in eq. (4) matches that of the autosomes before equality of the female and male autosomal means is applied [32, eq. 18]. Whereas female and male autosomal admixture random variables are identically distributed and their expectations can be written with a generic *δ*, the corresponding random variables differ for X chromosomes, and the coupled recursion in eqs. (3) and (4) cannot be quickly reduced to one equation. The mean X-chromosomal admixture in males (eq. (4)), however, has a similar form to the autosomal mean (eq. (5)).

We use the general mechanistic model presented here to derive closed-form expressions for the expected X-chromosomal admixture under two specific models of admixture, a single admixture event and constant admixture over time (Results).

### Data analysis: estimating admixture in African Americans

Many admixture studies have reported X-chromosomal and autosomal admixture estimates for African Americans (Table 2). For illustration, we focus on data from Cheng et al. [34], who provided one of the largest samples, reporting estimates of quantities close to those that appear in our model. Cheng et al. [34] estimated admixture fractions of the X chromosome and autosomes for 15,280 African-Americans from 14 studies.

**Table 2:** Studies that published estimates of the X-chromosomal and autosomal admixture fractions of an African American population. As the X chromosomes of females and males were considered jointly in the studies, we list the fraction of the sample that was female. In each study, the X chromosome shows elevated African ancestry compared to the autosomes. Values in the table were compiled from quantities reported in the studies, or by taking 1 minus an African estimate to obtain a European estimate, or vice versa, with the following exceptions. For Bryc et al. [31], the “autosome” estimates are genome-wide, including the X chromosome; the percentage of females was kindly provided by K. Bryc (pers. comm.). For Bryc et al. [23], we obtained an African X-chromosomal estimate by visual inspection of their Figure 4g, and we counted the number of females from their supplementary material. For Tian et al. [62], we computed the values from information reported in their Figure 5 caption.

| Reference | Mean ancestry estimates (%) |  |  |  | Percent female |
| --- | --- | --- | --- | --- | --- |
| | African<br>autosome<br>($E[H_{1,g}^A]$) | European<br>autosome<br>($E[H_{2,g}^A]$) | African X<br>chromosome<br>($E[H_{1,g}^X]$) | European X<br>chromosome<br>($E[H_{2,g}^X]$) | |
| **Cheng et al. 2009**<sup>34</sup> | 80.7 | 19.3 | 84.5 | 15.5 | 55.7 |
| **Bryc et al. 2015**<sup>31</sup> | 73.2 | 24.0 | 76.9 | 19.8 | 52.3 |
| **Bryc et al. 2010a**<sup>23</sup> | 77 | 23 | 81 | 19 | 66.3 |
| **Lind et al. 2007**<sup>19</sup> | 80 | 20 | 87.9 | 12.1 | 0 |
| **Tian et al. 2006**<sup>62</sup> | 81.8 | 18.2 | 87.1 | 12.9 | Not reported |

The mean admixture on the X chromosome reported by Cheng et al. [34] combines females and males into a single estimate, whereas we consider separate quantities for X-chromosomal admixture in females and males. In our notation, the mean X-chromosomal admixture across all individuals, 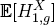, is a weighted average of admixture in females and males that takes into account the fraction of the sample that is female. Thus, for a sample divided into females and males in proportions *p*_*f*_ and *p*_*m*_ = 1 − *p*_*f*_, respectively, we can write

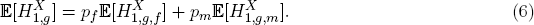

To compare admixture estimates from data to the mean admixture across individuals produced under mechanistic models, we compute the Euclidean distance between predictions for 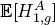, 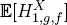, and 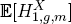 and observed values in the data, weighting the predicted female and male X-chromosomal admixture by eq. (6). Denoting by *q*_*A*_ and *q*_*X*_ the observed mean admixture in a specific source group for autosomes and the X chromosome, we evaluate

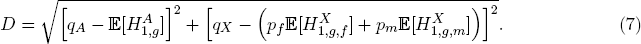

This quantity *D* measures the fit of a prediction under a model to the actual estimates of X-chromosomal and autosomal admixture for a data set that reports three quantities: estimates of admixture for a specific source population for autosomes and for the X chromosome, and the fraction of the admixed sample consisting of females.

## Results

To analyze the properties of X-chromosomal admixture in relation to the female and male contributions of two source populations, we consider two specific versions of our general admixture model. We then reinterpret African-American admixture estimates in light of these scenarios.

### A single admixture event

First, we study a special case in which no further contributions from the source population occur after the admixed population is founded in generation *g* = 1. In this scenario of a single admixture event [35], 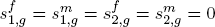, and 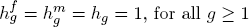, for all *g* ≥ 1.

Applying eqs. (3) and (4), we have for *g* = 1,

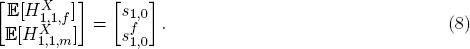

For *g* ≥ 2,

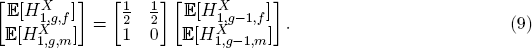

For *g* ≥ 2, the expected admixture fraction for a male X chromosome is simply the expected admixture fraction for a female X chromosome from the previous generation, or 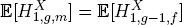. We can use this identity between the expected female admixture in generation *g* − 1 and male admixture in generation *g* to simplify the system of equations. For *g* ≥ 3, we then have

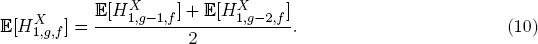

Denoting by *y*_*g*_ the quantity 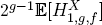, using eq. (10), for *g* ≥ 3, we have

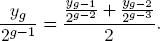

Multiplying both sides by 2^*g−*1^, we have a recursion *y*_*g*_ = *y*_*g*−1_ +2y_*g*−2_, with *y*_1_ = *s*_1,0_ and 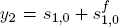. Then for *g* ≥ 3, *y*_*g*_ can be written 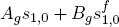, where *A*_*g*_ and *B*_*g*_ satisfy

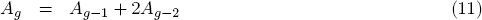

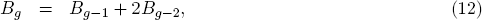

with *A*_1_ = 1, *A*_2_ = 1, *B*_1_ = 0, and *B*_2_ = 1. Noting that *B*_3_ = 1, we immediately observe that for *g* ≥ 3, *B*_*g*_ = *A*_*g*−1_, so that 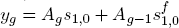. Then

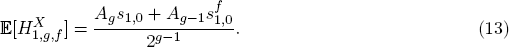

The recursion for *A*_*g*_ in eq. (11) with the initial conditions *A*_0_ = 0 and *A*_1_ = 1 gives the recursion for the Jacobsthal numbers, for which the closed form is *A*_*g*_ = [2^*g*^ − (−1)^*g*^]/3 (Online Encyclopedia of Integer Sequences A001045 website). We then have for *g* ≥ 1,

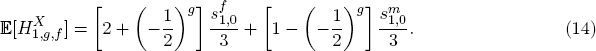

Using eq. (9), for *g* ≥ 2,

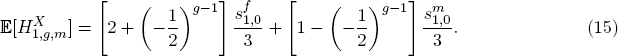

For the case of a single admixture event, with no further gene flow from the source populations to the admixed population, we use eqs. (14) and (15) to understand the behavior over time of the mean admixture on the X chromosome. Notably, unlike for the autosomes under a single-admixture scenario [32], the mean X-chromosomal admixture depends on the sex-specific contributions from the source populations, as well as on the time since admixture. With no sex bias 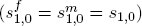, the mean X-chromosomal admixture is the same as the autosomal admixture, which is constant in time, equaling simply *s*_1,0_, the total contribution from *S*_1_.

Eqs. (14) and (15) oscillate over time (Fig. 2), as they incorporate a negative fraction raised to a power. The long-term limit of the X-chromosomal admixture fraction is

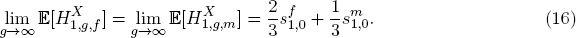

**Figure 2:**
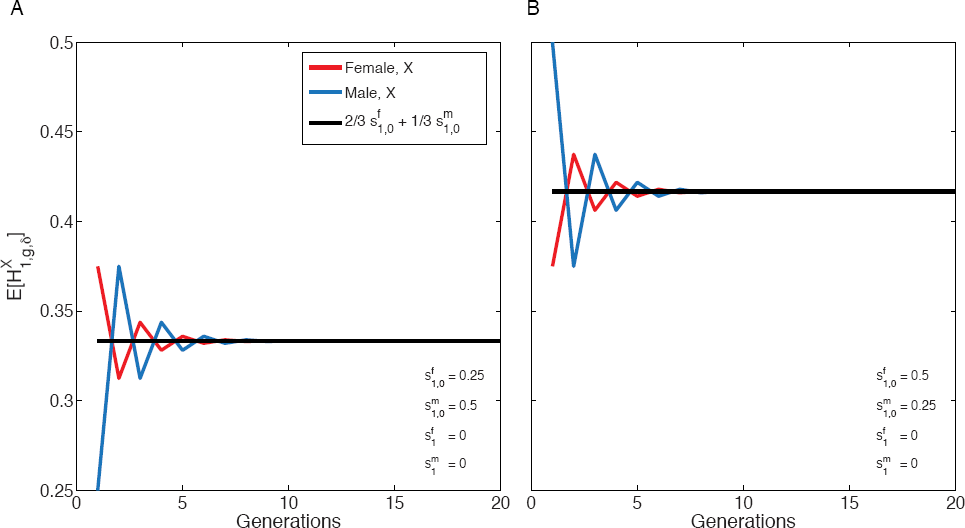
The mean of the X-chromosomal admixture fraction over time in females (red) and males (blue) from the admixed population for a single admixture event (eqs. (14) and (15)). The mean X-chromosomal admixture oscillates in approacing its limit (eq. (16)). The limiting mean, shown in black, is the same for X chromosomes in both females and males. (A) 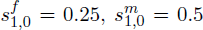. (B) 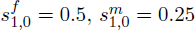.

Because 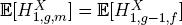 for all *g* ≥ 2, the expected admixture fraction sampled in males approaches the same limit over time as that sampled in females. The limit of the mean X-chromosomal admixture fraction fits the 2:1 ratio by which the X chromosome is inherited, following eq. (1), with the variables in eq. (1) viewed as the initial admixture values 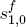 and 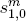. Thus, the limiting admixture, but not the transient admixture, matches the simple linear combination.

For recent admixture, we can calculate the difference between the expected admixture under the model and the limiting value. For *g* ≥ 1, we have

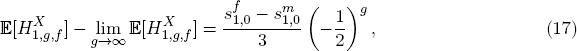

and for *g* ≥ 2,

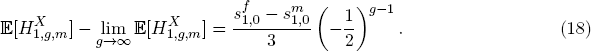

The differences in eqs. (17) and (18) provide a measure of the difference of a transient single-admixture model from the simpler linear combination in eq. (1), which is constant in time and does not consider differences between the mean admixture in females and males from the admixed population. The *g* → ∞ limit in eq. (16) agrees with eq. (1), but for small *g*, the differences in eqs. (17) and (18) can be as large as 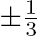, decreasing by a factor of 2 each generation. For a fixed *g*, the maximal absolute difference occurs when the sex bias is largest, that is, when 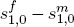 is 1 or −1. At the other extreme, with no sex bias and 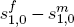, the simple linear combination exactly describes the X-chromosomal admixture throughout the history of the admixed population.

For six values of *g*, Figure 3 plots eq. (17) as a function of the difference between female and male contributions from 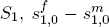. With no sex bias, 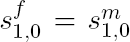, eq. (17) is zero, and our model follows eq. (1). Additionally, as *g* increases, the difference between our time-dependent model and eq. (1) becomes smaller, as eq. (1) gives the limiting behavior of the single-admixture model.

**Figure 3:**
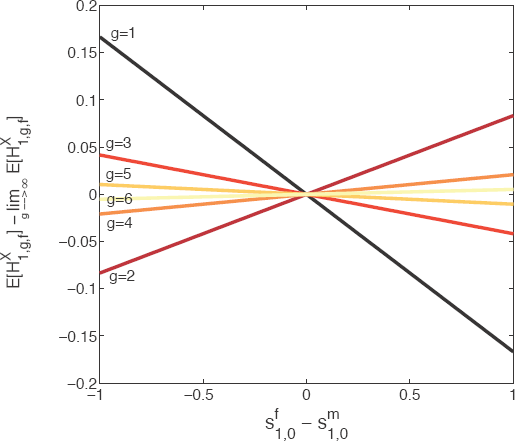
The difference between the expectation of the female X-chromosomal admixture fraction and its limit for *g* ∈ [1, 6] as a function of the difference between female and male contributions from *S*_1_ for a single admixture event (eq. (17)). As *g* → ∞, this quantity approaches zero. For small *g*, however, the limit 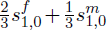 (eq. (16)) is a poor approximation. The difference oscillates so that the slope of the line is negative when *g* is odd and positive when *g* is even.

### Constant, non-zero admixture over time

Next, we consider the special case of constant, non-zero contributions from the source populations to the admixed population over time. As in Goldberg et al. [32], we can rewrite the sex-specific parameters as constants, 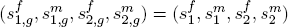, for all *g* ≥ 1. We maintain separate parameters for the founding contributions, 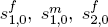, and 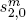. In this setting, *h*_*f*_ and *h*_*m*_ cannot both be 1, and at least one among 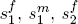, and 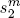 must be nonzero.

Using a generating function approach, we derive a closed-form solution for the mean X-chromosomal admixture fractions, 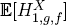 and 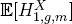. The mean depends on the number of generations of constant admixture, *g*, the initial conditions 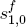 and 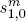, and the sex-specific contributions from the two source populations, 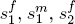, and 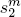. In the Appendix, we show that for each *g* ≥ 1,

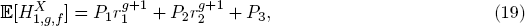

where *P*_1_, *P*_2_, and *P*_3_ are defined in eqs. (36)–(38), and *r*_1_ and *r*_2_ in eqs. (34) and (35). Using eqs. (27) and (19), we can write an expression for the male X-chromosomal admixture fraction,

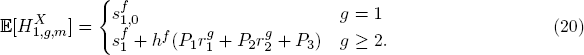

The limits of the mean X-chromosomal admixture fractions in eqs. (19) and (20) are

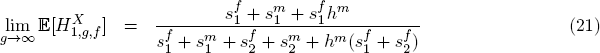

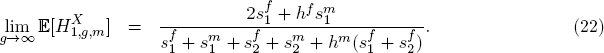

Unlike in the case of a single admixture event, the limit over time of the mean fraction of admixture from *S*_1_ for a constant admixture process is not a simple 2:1 weighting of the female and male contributions from *S*_1_. Notably, recalling 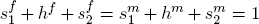, the limiting admixture fraction from *S*_1_ depends on the contributions both from *S*_1_ and from *S*_2_. The mean X-chromosomal admixture depends on the sex-specific contributions and cannot be reduced in terms of *s*_1_ and *s*_2_ only, as was possible for the autosomes [32, eq. 37].

For the autosomes, the limiting mean is the ratio of contributions from *S*_1_ to the total contributions from *S*_1_ and *S*_2_ combined [32, eq. 37], 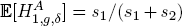. As was observed for the autosomes, the limiting mean X-chromosomal admixture in females can be viewed as the fractional contribution of X chromosomes from *S*_1_ in relation to the total number of X chromosomes from both source populations. Therefore, in eq. (21), the numerator has the same form as the denominator, incorporating only contributions from *S*_1_. X chromosomes from *S*_1_ present in a population of females from the admixed population come from one of three origins in the limit, identified by the three terms of the numerator. They can be directly contributed from *S*_1_ females or males, giving the terms 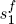 and 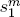. Alternatively, they can be contributed from admixed males who in turn received them from *S*_1_ females, producing the term 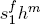; viewed in the limit, contributions from admixed females are already subsumed in the 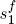 and 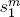 terms. Similar reasoning can be used to understand the limiting male admixture fraction, noting that the limiting mean for males is the sum of two quantities, the product of the limiting female mean and the fraction of females from the admixed population, *h*^*f*^, and the new influx of female contributions from *S*_1_, or 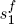 (eq. (27)).

The limits over time of the means in eqs. (21) and (22) depend only on the continuing contributions, not on the founding parameters. With no sex bias, 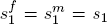 and *h*^*f*^ = *h*^*m*^ = *h*, and we can simplify the limiting mean female and male X-chromosomal admixture fractions:

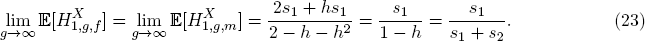

This limit is equivalent to the limiting mean of autsomal admixture from Verdu & Rosenberg [33, eq. 31]. That is, with no sex bias, the limiting X-chromosomal mean matches the limiting autosomal mean, 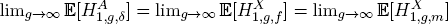.

Figure 4 plots the expectations of the X-chromosomal and autosomal admixture fractions over time for two scenarios with the same difference in female and male contributions, but different directions of sex bias. In Figure 4A, more males enter from *S*_1_ and more females from *S*_2_, leading to a lower mean admixture on the X chromosome than on the autosomes, with the mean in males smaller than the mean in females. Conversely, in Figure 4B, more females enter from *S*_1_ and more males from *S*_2_, leading to a larger mean for X chromosomes than for autosomes. The parameters are set so that for both panels, autosomal admixture is constant in time at 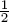 [32, eq. 37]. In both cases, the limiting mean X-chromosomal admixture in females lies intermediate between the mean autosomal admixture and the mean X-chromosomal admixture in males.

**Figure 4:**
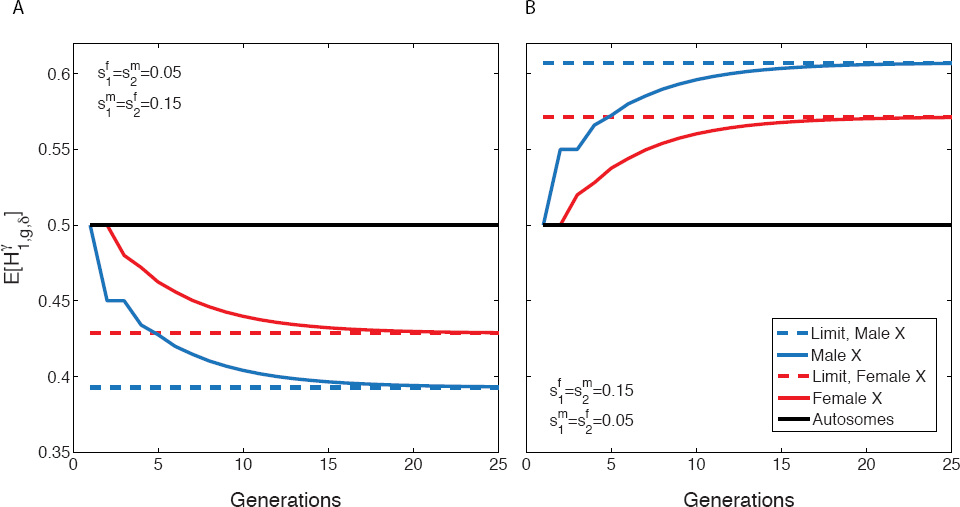
The expectation of the mean X-chromosomal and autosomal admixture fractions over time, with their associated limits, for constant ongoing admixture. (A) Male-biased admixture from population *S*_1_. (B) Female-biased admixture from population *S*_1_. The initial condition is 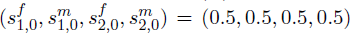. The autosomal admixture is constant over time because 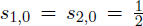 and *s*_1_ = *s*_2_; it is the same in both panels because it does not depend on the sex-specific contributions. The X-chromosomal admixture is different in females and males; it is smaller than the autosomal mean for male biased-admixture from population 1, and larger for female-biased admixture. The X-chromosomal mean is plotted using eqs. (19)-(22). The autosomal mean uses eq. 37 from Goldberg et al. [32].

Figure 5 plots the mean X-chromosomal admixture fraction for females and males (eqs. (19) and (20)) with the mean autosomal admixture fraction when both source populations have a female bias in contributions, 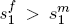 and 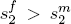. Though *S*_1_ has an excess of females, the mean X-chromosomal admixture is less than the autosomal admixture. Whereas an excess of females from a given source population might be viewed as generating elevated X-chromosomal admixture, when both populations have a sex bias in the same direction, the source population with the larger sex bias dominates the signal. It is tempting to interpret lower X-chromosomal than autosomal admixture as a signal of male-biased admixture from *S*_1_ and female-biased admixture from *S*_2_, but Figure 5 plots an example where admixture is female-biased in both source populations, and the simple interpretation of opposite biases in the two source populations is incorrect. A similar example could also have been produced with male bias in both source populations.

**Figure 5:**
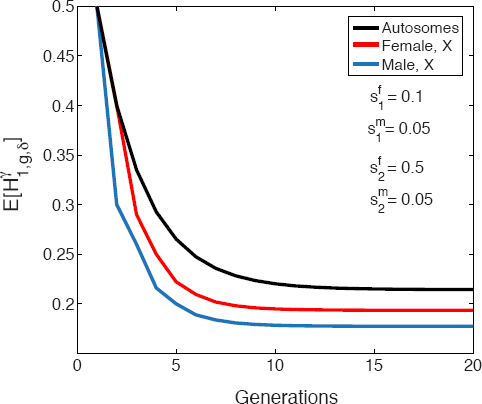
The expectation of the mean X-chromosomal and autosomal admixture fractions over time for constant admixture, with female-biased contributions from both source populations. The initial condition is 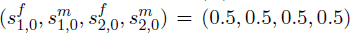. The mean X-chromosomal admixture 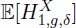, is smaller than the mean autosomal admixture, 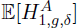, even though *S*_1_ has an excess of females, 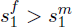. The expectation of X-chromosomal admixture is plotted using eqs. (19) and (20). The autosomal mean uses eq. 37 from Goldberg et al. [32].

We note that in the limit as the parameters 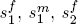, and 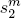 simultaneously approach zero, the constant-admixture model approaches the single-event model. Thus, we expect the limiting mean female and male X-chromosomal admixture fractions in the constant-admixture model to approach the corresponding limits in the single-event model. Indeed, by taking the limits of eqs. 19 and 20 as 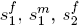 and 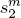 approach 0, we obtain eqs. 14 and 15, respectively. Interestingly, as we expect the trajectories of the X-chromosomal admixture fractions under constant admixture to approach corresponding single-event trajectories, we also expect the oscillatory pattern of the single-event model to occur in the constant admixture model for sufficiently small 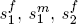, and 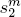. Indeed, we can find instances of oscillation around the limit in the constant-admixture model as well as in the single-admixture model, for example, for 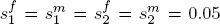 and initial condition 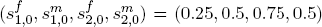 (Figure 6). As the continuing contributions in this case have no sex bias, the mean female and male X-chromosomal admixture and the mean autosomal admixture tend to the same limit.

**Figure 6:**
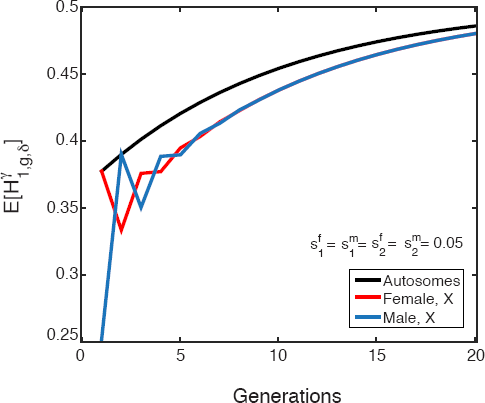
The expectation of the mean X-chromosomal and autosomal admixture fractions over time for constant admixture, in a case with small continuing contributions. The initial condition is 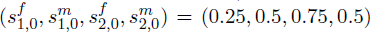. With sex bias in the founding generation and small continuing contributions, the mean X-chromosomal admixture has a pattern resembling the oscilating behavior seen for a single admixture event. The expectation of X-chromosomal admixture is plotted using eqs. (19) and (20). The autosomal mean uses eq. 37 from Goldberg et al. [32].

### Sex-biased admixture in African Americans

Studies of the genetic admixture history of African-American populations have consistently reported evidence for male-biased gene flow from Europe [15, 16, 19, 22, 23, 31]. Lind et al. [19] and Bryc et al. [31] estimated female and male contributions from Africa and Europe using mean admixture estimates for the X chromosome and autosomes. Both analyses followed the linear combination method of eq. (1), in which the mean autosomal admixture averages the female and male contributions, while the mean X-chromosomal admixture weights the female and male contributions by 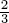 and 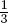, respectively. In our notation, the framework can be written

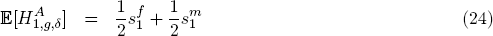

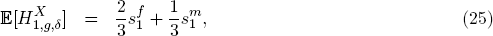

With 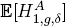 and 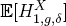 estimated from data.

We have demonstrated, however, that in autosomal admixture models, eq. (24) suggests a single admixture event [32], and in X-chromosomal admixture models, eq. (25) suggests the limit of a single admixture process (replacing parameters for the continuing contributions in these equations by corresponding parameters for admixture in the initial generation, 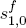 and 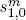, as in eq. (16)). For recent admixture in the single-admixture model, or in a model with continuing admixture after the founding of the admixed population, the mean admixture fractions from *S*_1_ for the X chromosome depend on the sex-specific contributions in a more complex way, incorporating the sex-specific contributions from *S*_2_ (eqs. (14), (15), (19)–(22)). Use of eqs. (24) and (25) implies consideration of a single-admixture model in its temporal limit, or otherwise is unsuited to the single-admixture and continuing admixture scenarios.

We used our refined predictions about 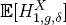 to estimate the sex-specific contributions for the African-American population from Cheng et al. [34] under two different models, a single admixture event with various times since admixture, and a constant admixture process with *g* = 15. Although both models underestimate the spatial and temporal complexity of the true admixture history of African Americans, these approximations enable us to illustrate the way in which estimates of the sex bias depend on assumptions about the admixture model.

The exact timing of the onset of significant admixture in the African-American admixture process is unknown, but assuming a generation time of 20-27 years, the Trans-Atlantic slave trade to North America can be regarded as having begun 14-20 generations ago [36]. As the mean X-chromosomal admixture fraction is near its limit for *g* in this range, both in the single-admixture model and in the constant-admixture model, the specific *g* ∈ [14, 20] only minimally changes the results. For our constant-admixture analysis, we chose a single value *g* = 15 for consistency.

***Single admixture event:*** Using the mean X-chromosomal and autosomal admixture reported by Cheng et al. [34] (Table 2), we calculated the sex-specific contributions 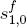 and 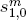 for specified values of *g* ∈ [2, 5], and in the *g* → ∞ limit. As Cheng et al. [34] reported admixture in a combined sample of females and males, we used eq. (6) to calculate the X-chromosomal admixture fraction 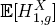 as a function of *p*_*f*_, *p*_*m*_, 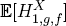, and 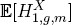. Writing 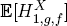 and 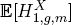 as functions of *g*, 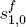, and 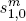 (eqs. (14) and (15)) then generated an equation for 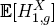 in terms of *p*_*f*_, *p*_*m*_, *g*, 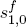, and 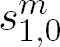, into which we inserted the reported values *qX* = 0.845 for 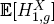 and (*p*_*f*_, *p*_*m*_) = (0.557, 0.443). We obtained a second equation for the autosomal admixture fraction from Goldberg et al. [32, eq. 34], 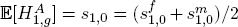, using *qA* = 0.807 for the empirical observation of 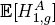. We then solved the pair of equations for 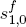 and 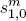, with *g* fixed.

For small *g* (2, 3, 4, 5) as well as in the *g* → ∞ limit, 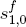, representing the contribution of females from the African source population, exceeds 0.9, and 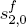, representing the female European contribution, is below 0.1 (Table 3). For males, the African contribution 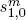 is ~0.7, whereas the European contribution 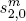 is ~0.3. Because of the oscillation of the mean X-chromosomal admixture (Fig. 2), the estimated ratio of male to female contributions oscillates around the limiting values. For contributions from Europe, in the *g* → ∞ limit we estimate 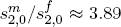. That is, in the temporal limit of a model of a single admixture event, for each female from Europe, ~3.89 males contributed to the gene pool of African Americans. The direction of this ratio is reversed for African contributions, with ~1.33 females for every African male. If we assume that the admixture was recent, for example *g* ≤ 5, then these values do vary substantially, with larger deviations from the limiting value of 3.89 occurring under more recent admixture.

**Table 3:** Estimates of the sex-specific contributions from Africa and Europe to African Americans, based on the Cheng et al. [34] data, inferred under our model for a single admixture event. The estimates depend on the time since admixture. All estimates show male-biased contributions from Europe and female-based contributions from Africa.

| Generations<br>since admixture | African contributions |  |  | European contributions |  |  |
| --- | --- | --- | --- | --- | --- | --- |
| | $s_{1,0}^f$ | $s_{1,0}^m$ | $s_{1,0}^f/s_{1,0}^m$ | $s_{2,0}^f$ | $s_{2,0}^m$ | $s_{2,0}^m/s_{2,0}^f$ |
| **2** | 0.943 | 0.671 | 1.40 | 0.057 | 0.329 | 5.77 |
| **3** | 0.912 | 0.702 | 1.30 | 0.088 | 0.298 | 3.39 |
| **4** | 0.926 | 0.688 | 1.35 | 0.074 | 0.312 | 4.22 |
| **5** | 0.919 | 0.695 | 1.32 | 0.081 | 0.305 | 3.77 |
| ∞ | 0.921 | 0.693 | 1.33 | 0.079 | 0.307 | 3.89 |

***Constant admixture over time:*** Assuming *g* = 15, we computed the mean female and male X-chromosomal admixture (eqs. (19) and (20)), and mean autosomal admixture given the four sex-specific contributions (eq. (5)), on a grid of possible parameter values, 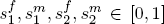 using 0.01 increments. We fixed the initial contributions 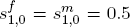, though by *g* = 15 the mean nears its limit, erasing the signal of the initial conditions (eqs. (21) and (22)). We calculated the distance *D* between the Cheng et al. [34] data and the computed values (eq. (7)).

The parameters 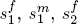, and 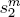 are not uniquely identifiable from the data, as we continue to have two equations, one describing autosomal admixture and one for X-chromosomal admixture, but now we consider four unknowns. We therefore examined sets of parameter values that produced *D* ≤ 0.01. Distance cutoffs of 0.001 and 0.1 gave rise to similar ranges for each parameter.

Figure 7 plots the set of parameter values that generate *D* ≤ 0.01. Because the female and male contributions separately sum to one, the possible range of each set of contributions is represented by a unit simplex. Figure 7A plots the marginal distributions of the set of parameter values with *D* ≤ 0.01. Contributions from *S*_1_ (Africans) vary over most of the permissible parameter space, but the *S*_2_ contributions (Europeans) take their values from a narrower range. Figure 7B plots *D* on the space of female contributions, 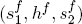, for values of 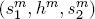 fixed at the median of the set of parameter vectors that produce 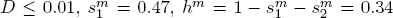, and 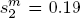. Figure 7C plots *D* as a function of the male contributions, 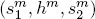, with female contributions fixed at the median from the distribution of parameter values for which *D* ≤ 0.01, 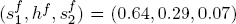. In both panels, the parameter sets closest to the data in terms of *D* permit a large range of possible contributions from *S*_1_, Africans, but only a smaller range of possible contributions, both female and male, from Europeans, *S*_2_.

**Figure 7:**
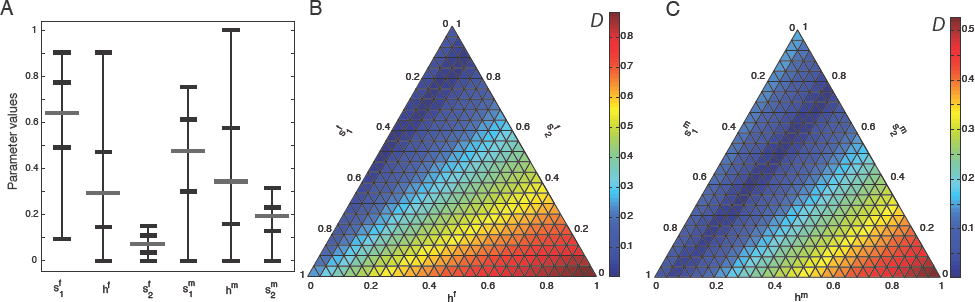
Sex-specific contributions estimated from the data of Cheng et al. [34]. (A) The range, median, and 25th and 75th percentiles of the sets of sex-specific contributions for which the Euclidean distance *D* (eq. (7)) between model-predicted admixture and observed admixture from Cheng et al. [34], was at most 0.01. The range of values for 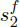 and 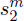, the contributions representing Europeans (*S*_2_), is smaller than those representing Africans (*S*_1_). (B) Female contributions as a function of *D*. (C) Male contributions as a function of *D*. For (B), the male contributions are fixed at their median values producing *D* ≤ 0.01, 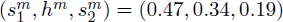. For (C), the female contributions are fixed in a corresponding manner, at 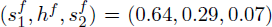. The space of possible values for 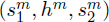 or 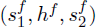 is the unit simplex. Parameter values were tested at increments of 0.01 for each quantity.

Figure 8A plots the natural logarithm of the ratio of male to female contributions for parameter sets with *D* ≤ 0.01. With no sex bias, this quantity is zero, with more males than females it is positive, and with more females than males it is negative. Most points plotted from *S*_1_, Africans, are female-biased, whereas contributions from *S*_2_, Europeans, are predominantly male-biased. The median ratio of males to females from Europe is 2.67 males per female, compared to a median 1.32 females per male from Africa. Note, however, that these patterns do not hold for all parameter sets. For 26.15% of values with *D* ≤ 0.01, more male than female contributions occur from Africa. For 9.25% of values with *D* ≤ 0.01, more female than male contributions occur from Europe.

**Figure 8:**
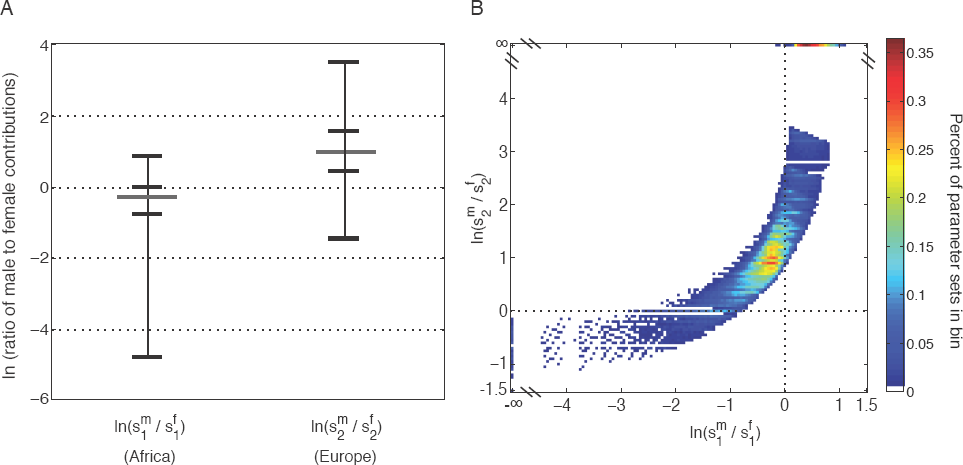
The natual logarithm of the ratio of male to female contributions in African Americans, as inferred from the data of Cheng et al. [34]. (A) The range (excluding infinity, produced when a parameter value is zero), median, and 25th and 75th percentiles of the natural logarithm of the ratio of male to female contributions from *S*_1_ (Africans) and *S*_2_ (Europeans) separately for the sex-specific contributions that produced *D* ≤ 0.01 (eq. (7)). Values from Africans, *S*_1_, are largely below 0, or female-biased, whereas contributions from Europeans, *S*_2_, are mostly above 0 and male-biased. Approximately 26.15% of the contributions from Africans are male-biased and 9.25% from Europeans are female-biased. This pattern is typically observed when a still larger sex bias occurs in the other population. (B) The logarithm of the ratio of male and female contributions from *S*_2_ on the y-axis and the corresponding ratio for *S*_1_ on the x-axis, plotted by the density of points in 0.05 square bins. For the cases with male bias in Africa 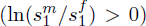, the level of male bias in Europe is also positive; for the cases with female bias in Europe 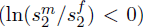, the level of female bias from Africa is also negative. Parameter sets in which at least one parameter is 0, and therefore, we have values of +∞ or −∞ for ln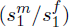 or ln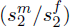, appear in bins on the edge of the plot for convenience. These bins contain a substantial number of parameter sets.

Although the pattern is initially surprising, scenarios of female bias from Europe or male bias from Africa accord with the phenomenon depicted in Figure 5. In the former scenario, both populations have an excess of females, and the strong female bias in the contributions from Africa can overwhelm the number of European X chromosomes in the gene pool of African Americans, producing greater African admixture on the X chromosome than on autosomes. This effect is demonstrated in Figure 8B, which plots the logarithm of the ratio of male to female contributions from *S*_2_ against the corresponding quantity from *S*_1_. The ratios of male to female contributions in *S*_2_ on the y-axis and *S*_1_ on the x-axis are correlated. The values that are negative on the y-axis, indicating a female bias from *S*_2_, also show the strongest female bias from *S*_1_, that is, the most negative 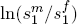 on the x-axis (an additional 2.34% of parameter sets show no sex bias in the European population; the female and male contributions are equal and 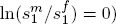. Analogously, the strongest male bias from Africa is also observed when the excess of males from Europe is largest (Fig. 7B).

## Discussion

Under a two-sex mechanistic admixture model with sex-biased admixture, we have demonstrated that the relationship between X-chromosomal and autosomal admixture fractions depends both on the time since admixture and on the model of admixture, and does not simply follow a prediction from the fractions of X chromosomes and autosomes present in females. Using the mechanistic framework, we have reinterpreted African American admixture values computed from a non-mechanistic perspective, estimating sex-specific parameters and levels of sex bias in African and European source populations. This analysis uncovers a counterintuitive case in which female-biased or male-biased contributions in the same direction occur both from Africans and Europeans in a manner consistent with estimated mean X-chromosomal and autosomal admixture fractions in African Americans, rather than an an African female bias and a European male bias.

### Estimating sex bias using X-chromosomal and autosomal admixture

Differences between X-chromosomal and autosomal admixture estimates are sometimes used to demonstrate the occurrence of sex bias, even without estimating sex-specific contributions [21, 23, 24, 26, 27, 37]. Higher estimated admixture from a population *S*_1_ for X chromosomes than for autosomes is taken as evidence of a female bias from *S*_1_ and a male bias from a second population *S*_2_. We have found, however, that the pattern is in principle compatible with two additional possibilities (Fig. 7): a female bias from both populations, with a larger female bias from *S*_1_, or a male bias from both populations, with a larger male bias from *S*_2_.

We considered a general mechanistic model of admixture, focusing specifically on models of a single admixture event and constant admixture. For a single admixture event, mean autosomal admixture over time is constant, depending only on the total contributions from the source populations, and not on the sex-specific contributions or time [32]. Mean X-chromosomal admixture, on the other hand, is variable over time—in such a way that given the number of generations *g* and observed levels of X-chromosomal and autosomal admixture, the initial sex-specific contributions from the two source populations are identifiable. Because of the oscillation of the mean X-chromosomal admixture over time, depending on the time since admixture, the sex bias in admixture contributions can be either overestimated or underestimated using the 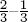 linear combination (eq. (1)).

For constant admixture, estimated mean X-chromosomal and autosomal admixture values do not uniquely identify the female and male contributions over time. Therefore, instead of point estimates for the sex-specific contributions and their ratio, we have reported a measure of the compatibility of parameter values and data over a range of values of the parameters (Figs. 6 and 7). It is possible, however, that by examining higher moments of the distribution of admixture estimates across individuals, the sex-specific contributions might become identifiable [32].

### Theoretical population genetics of the X chromosome

The complex signature of admixture we have observed for the X chromosome is reminiscent of other X-chromosomal phenomena in theoretical population genetics, including results related to effective population size, allele-frequency dynamics, and numbers of ancestors. For effective population size, *N*_*e*_, when female and male population sizes are equal, the ratio of X-chromosomal to autosomal values is 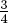 [38–40]. In the same way that various forms of sex difference between females and males—in such parameters as the number of individuals, the variance of reproductive success, and migration rates—cause the basic X-to-autosomal *N*_*e*_ ratio to differ from 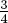 [1, 3, 4, 6, 8, 12–14, 39, 41], transient dynamics and ongoing admixture cause the X-chromosomal admixture fraction to differ from a linear combination of female and male contributions with coefficients 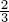 and 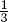.

The X-chromosomal admixture fraction in our single-admixture model also has similarities to the allele-frequency trajectory in the approach to Hardy-Weinberg equilibrium in a one-locus X-chromosomal model. In that model [42–44], the frequency of an allele on female X chromosomes depends on the frequency in both females and males in the previous generation, whereas the frequency on male X chromosomes depends only on the frequency in females. If an allele frequency differs between females and males at the first generation, then during the approach to equilibrium, the frequency in males matches the corresponding frequency in females of the previous generation; both the female and male frequencies oscillate around the same limit. The equilibrium frequency in turn has a 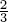 contribution from the initial female frequency and 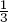 from the male frequency.

All these phenomena from the one-locus model—female values dependent on both female and male values from the previous generation, male values dependent only on the female value, males lagging one generation behind females, oscillations around the same limit, and a limiting linear combination of 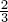 and 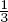—appear in our X-chromosomal admixture model with a single admixture event. An additional parallel lies in the difference between the X chromosome and the autosomes: in the one-locus model for the X chromosome, Hardy-Weinberg equilibrium is achieved in the limit, and for the autosomes, it is achieved in one generation. For admixture on the autosomes, the mean admixture is constant after the founding of the admixed population, and for the X chromosome, the mean X-chromosomal admixture approaches a limit rather than remaining constant in time.

Yet another connection to theoretical population genetics of the X chromosome comes from the form of the recursive sequence in eq. (13). The Jacobsthal sequence in the single-admixture model is obtained from a generalization of the Fibonacci sequence [45]. Both the Jacobsthal and Fibonacci sequences appeared in breeding system models as early as a century ago [42, 46–48], with the Jacobsthal sequence providing the numerators of allele frequencies at a sex-linked locus in a model that amounts mathematically to a special case of our single-event admixture model [42, 47]; the Fibonacci sequence is well-known as the number of genealogical ancestors of a haplodiploid system *g* generations ago [49]. Specifically, for the pair of X chromosomes in a female, the number of ancestors is entry *g* + 2 in the Fibonacci sequence *F*_*n*_, or equivalently, the sum of the numbers of maternal and paternal ancestors, *F*_*g*+1_ and *F*_*g*_, respectively. The number of genealogical ancestors for a male X chromosome in generation *g* is *F*_*g*+1_. Our recursion *A*_n_ = *A*_*n*−1_ + 2*A*_*n*−2_ (eq. (11)) generates a different sequence, but its form is similar to the Fibonacci recursion *F*_*n*_ = *F*_*n*−1_ + *F*_*n*−2_.

### Sex-biased admixture in African-Americans

Our analysis of sex-biased admixture in African Americans has similarities but a number of noteworthy differences from earlier nonmechanistic analyses. Under our single-admixture model, the estimate of ~4 European males for every European female accords with estimated ratios of 3 to 4 from previous studies [19, 31]. For even *g*, more African females and European males contribute than in the *g* → ∞ limit; odd *g* values produce the opposite pattern.

For the constant admixture model, the median estimated ratios of male to female contributions from Europe and Africa are ~2.67 males per female from Europe and ~1.32 females per male from Africa (Figs. 6 and 7). The estimated male bias in contributions from Europe is lower for a constant admixture history than for a single admixture event.

For both a single admixture event and constant admixture over time, the point estimates of the ratio of males to females have a larger male contribution from Europe and female contribution from Africa. In the single-admixture model, the excess African ancestry on the X chromosome compared to the autosomes implies a female-biased contribution from Africa and a male-biased contribution from Europe. For constant admixture, surprisingly, a sex bias in one source alone can produce the observed pattern without a sex bias in the other (Fig. 7). In fact, ~27% of the parameter sets most similar to the data (*D* ≤ 0.01) show no sex bias or have larger male than female contributions from Africa; ~12% show no sex bias or have larger female than male contributions from Europe. Note that in obtaining this result, we have given equal weight to all values of *D* below a cutoff, rather than giving more weight to parameter choices producing lower *D*; however, owing to the existence of ridges in the parameter space that have similarly small values of *D*, for different choices of the cutoff, similar results are produced.

An excess of African mitochondrial and European Y-chromosomal haplotypes has been described in African Americans [15, 16, 19, 22, 50]. Similar phenomena to those we observed for X chromosomes and autosomes could affect the Y–mtDNA comparisons: in other words, a strong excess of males from Europe compared to European females, even if more males than females contribute from African populations, could give rise to a larger fraction of European Y chromosomes in the African-American gene-pool without a female bias for the African source population. Such a process could also explain the excess African ancestry of mtDNA.

### Conclusions

We have presented a model to describe the effect of admixture on the X chromosome, deriving a theoretical framework that considers the impact of sex bias during the admixture process. The model can be used to understand the sex-specific contributions from source populations to an admixed population. We have found that because of model dependence, time dependence, and a lack of identifiability of admixture parameters from mean admixture alone, a variety of admixture processes and parameter values might be compatible with estimates of the mean admixture on X chromosomes and the autosomes. Factors we have not studied, including mutation, recombination, selection, and genetic drift can differentially affect the X chromosome and autosomes [10, 13, 41, 51–55], potentially further complicating the estimation of admixture parameters. Examination in the context of sex-biased models of more detailed summaries of admixture patterns, including higher moments of the admixture distributions [32, 33, 56] and distributions of the lengths of admixture tracts [56–58], will assist in refining the estimation of sex-biased admixture histories.

We demonstrated that female and male X-chromosomal admixture have different expectations under mechanistic admixture models. With the exception of studies restricted to males [19], however, past studies of sex-biased admixture have generally reported composite X-chromosomal admixture estimates only in pooled collections of females and males. We recommend that such studies report estimates separately in females and males. Even if the values are similar, the difference between X-chromosomal admixture estimates in females and males contains information about parameters of the admixture process. Autosomal admixture estimates obtained only in females and only in males are identically distributed [32], so that a pooled estimate is more sensible than in the X-chromosomal case; nevertheless, autosomal admixture estimates reported separately in females and males can enable an informative comparison with a corresponding pair of X-chromosomal values.

Curiously, we have found that African American admixture patterns do not necessarily imply a simultaneous African female bias and European male bias in the genetic ancestry of the population, and that in both source populations, it is in principle possible on the basis of genetic admixture patterns that both source populations had female bias or that both had male bias, albeit at different magnitudes. The latter interpretation, of male biases both in Europeans *and* in Africans is plausible in light of historical scholarship on demographic contributions to African Americans [36, 59–61], documenting both the well-known contributions of male European slave owners and asymmetric mating practices involving African females and European males, as well as an overrepresentation of males among African slaves arriving in North America. For less well-documented cases, our model highlights that if a difference in X-chromosomal and autosomal admixture is observed, it is important to consider the possibility that rather than opposite sex biases in the two populations, both populations might have the same type of sex bias.

## Acknowledgments.

We thank Carlos Bustamante, Filippo Disanto, Michael D. Edge, Marc Feldman, Ethan Jewett, Hua Tang, and Paul Verdu for useful discussions. We acknowledge support from NIH grant R01 HG005855 and a National Science Foundation Graduate Research Fellowship.

## Web Resources

On-Line Encyclopedia of Integer Sequences A001045 website, http://oeis.org/A001045.

## Appendix: Solving for 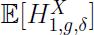 under constant admixture

Here, we obtain the closed form for the mean X-chromosomal admixture fraction in a random female and a random male from the admixed population in the special case of constant admixture over time. Using the constant sex-specific parameters, 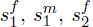, and 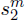, we rewrite the expectations in eq. (4). For *g* = 1, 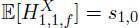, and for *g* ≥ 2, we have

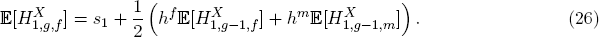

Similarly, sampling males in the admixed population, for *g* = 1, 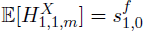. For *g* ≥ 2,

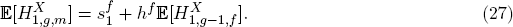

Using equations (26) and (27), we derive a generating function for the mean female X-chromosomal admixture fraction, which we then use to find closed-form expressions for 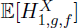 and 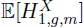. Because 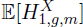 depends only on constants and 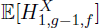, we first find an expression for 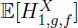 which we then use to report 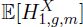.

First, using eq. (27), we can rewrite eq. (26) as a second-order recursion of a single variable,

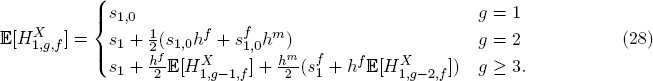

We simplify the notation by defining 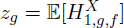. For *g* ≥ 3, we have,

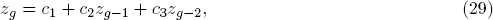

with 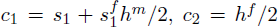, and *c*_3_ = *h*^*f*^ *h*^*m*^/2. Eq. (28) gives *z*_1_ = *s*_1,0_ and 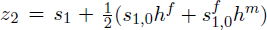.

Define a generating function 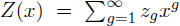 whose coefficients *z*_*g*_ represent the values of 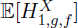 in each generation. As the admixed population does not yet exist in generation 0, 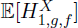 and *Z*(*x*) are undefined for *g* = 0. For convenience, we therefore work with *W*(*x*) = *Z*(*x*)/*x*, setting *w*_*g*_ = *z*_*g*+1_ for *g* ≥ 0. We then have

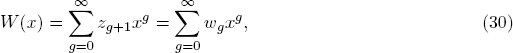

and *w*_*g*_ = *c*_1_ + *c*_2_*w*_*g*−1_ + *c*_3_*w*_*g*−2_ for *g* ≥ 2. Using eq. (30), it follows that

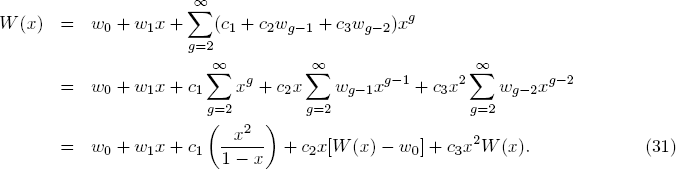

Solving for *W*(*x*), we have

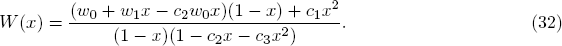

We can decompose the expression in eq. (32), producing

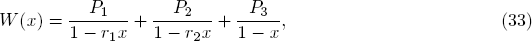

where *r*_1_ and *r*_2_ are reciprocals of the two roots of 1 − *c*_2_*x* − *c*_3_*x*_2_,

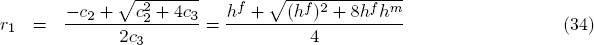

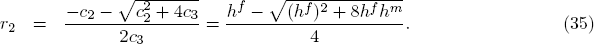

Setting eq. (32) equal to eq. (33), we have

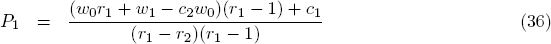

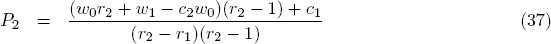

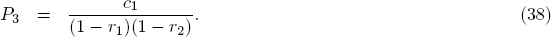

The Taylor expansion of eq. (33) around *x* = 0 then gives

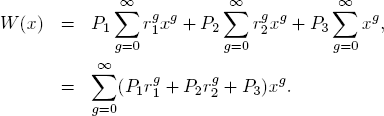

Therefore, for *g* ≥ 0, 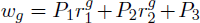, and the closed-form expression for the X-chromosomal female mean admixture fraction in generation *g* ≥ 1, 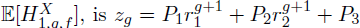. We report this result in the main text as eq. (19), using it to obtain 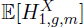 in eq. (20).

Because for *h*^*f*^, *h*^*m*^ ∈ [0, 1], *r*_1_ monotonically increases in *h*^*f*^ and *h*^*m*^ and *r*_2_ monotonically decreases, the maxima and minima of *r*_1_ and *r*_2_ occur at the boundaries of the closed interval [0, 1]. Using eqs. (34) and (35), we have *r*_1_ ∈ [0, 1) and 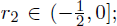 we exclude *r*_1_ = 1 and 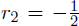, as *h*^*f*^ and *h*^*m*^ cannot both be 1. Because |*r*_1_|, |*r*_2_| < 1, the mean X-chromosomal admixture fractions in eqs. (19) and (20) approach limits as *g* → ∞. Using eqs. (19) and (20), we can find expressions for the limits of the mean of the X-chromosomal admixture fractions,

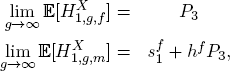

which can be simplified to give eqs. (21) and (22).

